# Metabolically activated macrophages in mammary adipose tissue link obesity to triple-negative breast cancer

**DOI:** 10.1101/370627

**Authors:** Payal Tiwari, Ariane Blank, Chang Cui, Kelly Q. Schoenfelt, Guolin Zhou, Yanfei Xu, Ajay M. Shah, Seema A. Khan, Marsha Rich Rosner, Lev Becker

## Abstract

Obesity is associated with increased incidence and severity of triple-negative breast cancer (TNBC); however, mechanisms underlying this relationship are incompletely understood. Macrophages, which accumulate in adipose tissue and are activated during obesity, are an attractive mechanistic link. Here, we show that, during obesity, murine and human mammary adipose tissue macrophages adopt a pro-inflammatory, metabolically-activated (MMe) macrophage phenotype that promotes TNBC stem-like markers and functions, including increased tumorsphere growth *in vitro* and tumor-initiating potential *in vivo*. We demonstrate that MMe macrophages release cytokines in an NADPH oxidase 2 (NOX2)-dependent manner that signal through glycoprotein 130 (GP130) on TNBC cells to promote their stem-like properties. Accordingly, deleting *Nox2* in myeloid cells or depleting GP130 in TNBC cells attenuates the ability of obesity to drive TNBC tumor formation. Our studies implicate MMe macrophage accumulation in mammary adipose tissue during obesity as a mechanism for promoting TNBC stemness and tumorigenesis.

**HIGHLIGHTS:**

⁘ Obesity promotes TNBC tumor formation and stemness.
⁘ Mammary adipose tissue macrophages are metabolically activated (MMe) in obese mice and humans.
⁘ MMe macrophages in mammary adipose tissue contribute to obesity-induced stemness.
⁘ MMe macrophages promote TNBC stemness through GP130 signaling.

## INTRODUCTION

Obesity is a major modifiable risk factor for breast cancer and is responsible for approximately 20% of cancer deaths (Calle et al., 2003). In addition to its role in breast cancer pathogenesis, obesity is recognized as a marker of poor prognosis in pre- and post-menopausal women with breast cancer (Chan and Norat, 2015). Epidemiological studies have linked obesity with increased risk of developing different subtypes of breast cancer, including triple-negative breast cancer (TNBC) (Vona-Davis et al., 2008; Trivers et al., 2009; Pierobon and Frankenfeld, 2013), a particularly aggressive form of breast cancer with poor outcome and few therapeutic options. Among TNBC patients, progression and disease-free survival (DFS) are strongly correlated with excessive weight and obesity (Choi et al., 2016). However, mechanisms by which obesity leads to a worse breast cancer prognosis are incompletely understood.

One clue to its action is that obesity causes chronic inflammation. A recent study showed that obesity-induced neutrophil accumulation in the lung can promote breast cancer metastasis (Quail et al., 2017). In addition to inflammation at metastatic sites, obesity also promotes local inflammation in adipose tissue which is mediated by macrophage infiltration and activation (Xu et al., 2003; Lumeng and Saltiel, 2011). Obesity-induced inflammation in mammary adipose tissue (Howe et al., 2013; Vaysse et al., 2017) may be of particular significance because breast cancers form in this niche and inflammation promotes stem-like properties in cancer cells and an increased propensity to form tumors (Grivennikov et al., 2010). Thus, pro-inflammatory macrophage accumulation in mammary fat may augment TNBC tumor formation during obesity.

Macrophages are heterogeneous and have been broadly classified as either classically (M1) or alternatively (M2) activated (Gordon and Taylor, 2005). Th2 mediators (e.g. IL-4) drive the M2 phenotype, which scavenges debris, and promotes angiogenesis and tumor growth (Noy and Pollard, 2014). In contrast, the M1 phenotype is promoted by Th1 mediators (e.g. LPS and IFNγ) and is characterized by the production of pro-inflammatory cytokines and anti-tumor activity (Pyonteck et al., 2013). Earlier studies showed that obesity promotes an M1-like ATM phenotype (Lumeng et al., 2007), which would be expected to oppose tumor formation. However, more recent studies from multiple labs have challenged the notion that obesity supports an M1 phenotype (Xu et al., 2013; Kratz et al., 2014).

Studies from our group showed that obesity produces a pro-inflammatory ‘metabolically activated’ (MMe) ATM phenotype that is both mechanistically and functionally distinct from the M1 phenotype (Kratz et al., 2014; Coats et al., 2017). Although we showed that MMe macrophages accumulate in visceral and subcutaneous adipose tissue of obese humans and mice, their presence in mammary fat, and their ability to promote TNBC tumor formation have not been explored.

Here, we show that MMe macrophages accumulate in mammary fat of obese mice and humans. We demonstrate that these MMe macrophages secrete cytokines in an NADPH oxidase 2 (NOX) dependent manner, that signal through glycoprotein 130 (GP130) on murine and human TNBC cells to promote stem-like properties and tumor formation during obesity. These findings reveal an important mechanism by which obesity enhances TNBC tumorigenesis.

## RESULTS

### Diet-induced obesity promotes TNBC stemness and tumor formation

To determine if diet-induced obesity (DIO) promotes TNBC tumorigenesis, we first studied genetically engineered C3(1)-TAg mice, which spontaneously develop TNBC-type tumors in multiple mammary glands (Green et al., 2000). Female C3(1)-TAg mice (on the FVB/N background) were fed a low-fat diet (LFD) or high-fat diet (HFD) for 12 weeks. Although FVB/N mice are somewhat protected from DIO (Montgomery et al., 2013), HFD-fed mice had increased body weight, fasting glucose, and mammary/visceral fat pad weight compared to LFD-fed mice (**Figs. 1A-C**).

**Fig. 1.**
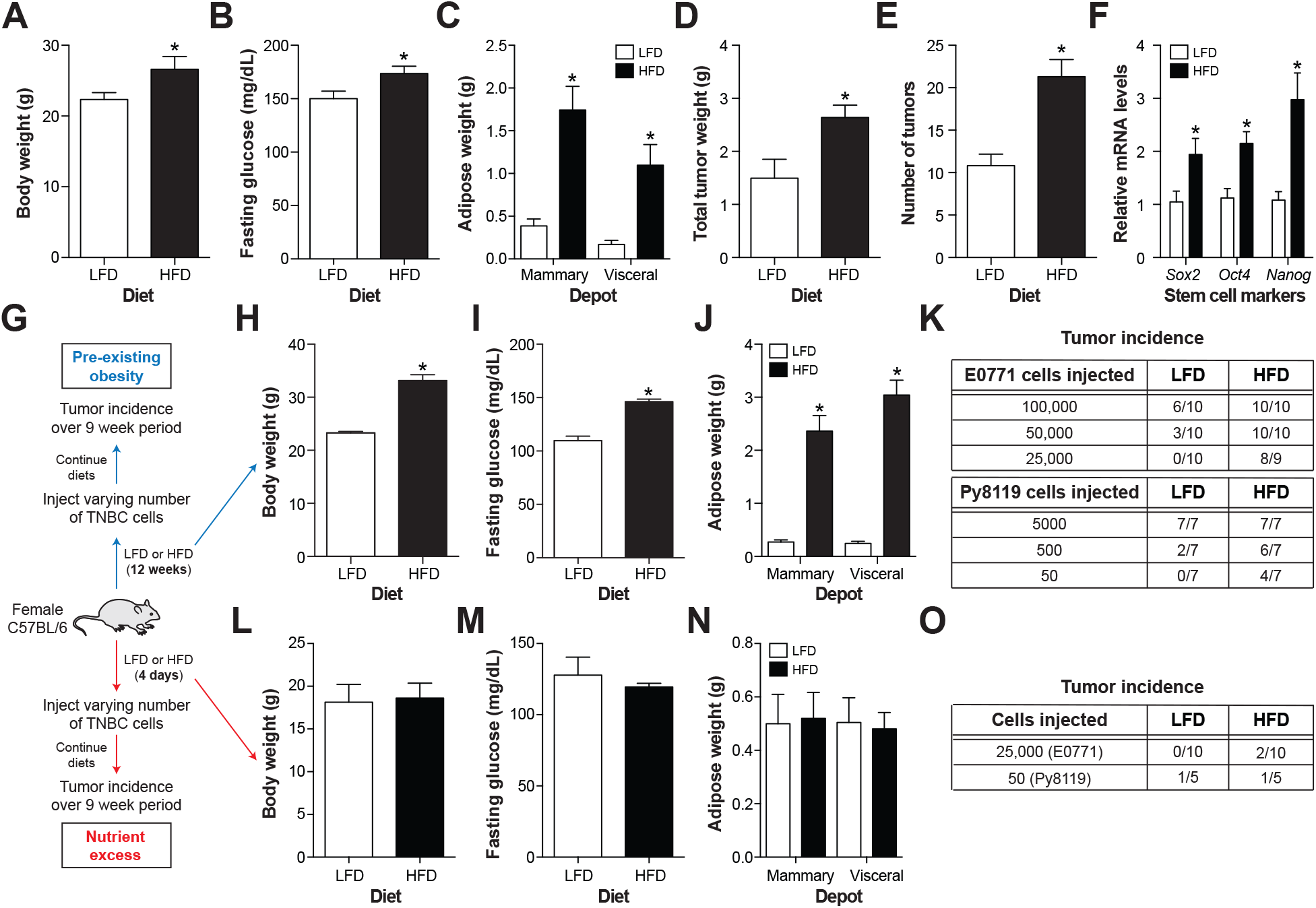
Diet-induced obesity promotes TNBC cell tumor formation. *<u>Panels A-F</u>*: Female C3(1)-TAg mice were fed a low-fat diet (LFD) or high-fat diet (HFD) for 12 weeks. *<u>Panel A</u>*: Body weight. *<u>Panel B</u>*: Fasting glucose levels. *<u>Panel C</u>*: Mammary and visceral adipose tissue weight. *<u>Panel D</u>*: Tumor weight. *<u>Panel E</u>*: Number of tumors/mouse. *<u>Panel F</u>*: Stem cell marker expression in isolated cancer cells. *<u>Panel G</u>*: Female C57BL/6 mice were fed a LFD or HFD for 12 weeks to create pre-existing obesity or 4 days to create nutrient excess. *<u>Panels H-K</u>*: Pre-existing obesity; Body weight (*<u>Panel H</u>*). Fasting glucose levels (*<u>Panel I</u>*). Mammary and visceral adipose tissue weight (*<u>Panel J</u>*). Limiting dilution assay of E0771 or Py8119 TNBC cells injected into mammary fat of mice (*<u>Panel K</u>*). *<u>Panels L-O</u>*: Nutrient excess; Body weight (*<u>Panel L</u>*). Fasting glucose levels (*<u>Panel M</u>*). Mammary and visceral adipose tissue weight (*<u>Panel N</u>*). Limiting dilution assay of E0771 or Py8119 TNBC cells injected into mammary fat of mice (*<u>Panel O</u>*). Results are mean ± SEM. *, *p*<0.05 Student’s *t*-test. n=5-10 mice/group.

As previously reported (Mustafi et al., 2017), DIO increased the total tumor burden in C3(1)-TAg mice (**Fig. 1D**), and this increased burden was due, in part, to the presence of more tumors in obese mice (**Fig. 1E**), suggesting that DIO may promote tumor initiation in a genetically engineered mouse model of TNBC.

The tumor initiating capacity of cancer cells has been linked to their stem-like properties (Nguyen et al., 2012). We therefore quantified the expression of the stem-associated genes *Sox2*, *Oct4*, and *Nanog* (Klonisch et al., 2008), in cancer cells isolated from tumors of lean and obese C3(1)-TAg mice. We found that obesity increased the mRNA expression of *Sox2*, *Oct4*, and *Nanog* in cancer cells (**Fig. 1F**), suggesting that obesity may create an environment that enhances stemness in TNBC cells.

To test this possibility, we switched to a syngeneic orthotopic transplant model in obesity-prone C57BL/6 mice. This model allowed us to establish an obese environment, and then determine whether this pre-existing environment is more capable of supporting the tumor-initiating potential of TNBC cells, a key function ascribed to cancer stem cells.

We fed female C57BL/6 mice a LFD or HFD for 12 weeks to induce obesity, hyperglycemia, and mammary and visceral fat expansion (**Figs. 1G-J**). At this time, murine E0771 or Py8119 TNBC cells were injected at limiting dilutions into mammary fat pads of lean or obese mice, diets were continued, and their tumor-initiating potential was assessed using a limiting dilution assay. We found that DIO decreased the number of E0771 or Py8119 cells required to form tumors (**Fig. 1K**) as well as tumor latency (**Fig. S1**), reinforcing the notion that obesity promotes TNBC tumor formation.

The ability of DIO to promote the tumor-initiating potential of TNBC cells could be due to high nutrient levels that support tumor growth, or due to a pathophysiological change induced by chronic high-fat feeding. To differentiate between these possibilities, we fed female C57BL/6 mice for 4 days with a LFD or HFD to create conditions of nutrient excess without changes in body weight, fasting glucose, or mammary/visceral adipose tissue mass (**Figs. 1G, 1L-N**). At this time, we injected E0771 or Py8119 cells, continued diet feeding, and quantified tumor incidence over a 7-week period.

Short-term pre-exposure to HFD did not support increased E0771and Py8119 tumor formation (**Fig. 1O**), even though mice were maintained on the HFD for 7 weeks following tumor cell injection. These data suggest that the induction of TNBC tumor-initiating potential requires a state of pre-existing obesity, and is not simply reliant on nutrient excess.

### Obese mammary ATMs induce stem-like properties in TNBC cells

Obesity has been characterized as a chronic inflammatory state (Lumeng and Saltiel, 2011), and pro-inflammatory cytokines have been shown to enhance the tumor-initiating potential of many types of cancer cells (Nguyen et al., 2012). Since ATMs are an important source of cytokines during obesity, we hypothesized that an increase in pro-inflammatory ATMs in mammary fat might help to explain how DIO promotes TNBC tumor formation.

To begin to test this hypothesis, we determined whether DIO could increase the number of mammary fat ATMs and/or their pro-inflammatory cytokine expression in female C57BL/6 mice fed the HFD for 12 weeks. Although DIO substantially increased mammary adipose tissue mass (see Fig. 1C), the percent CD45+ immune cells or ATMs in the stromal vascular cells (SVC) were not increased (**Fig. 2A, Fig. S2**).

**Fig. 2.**
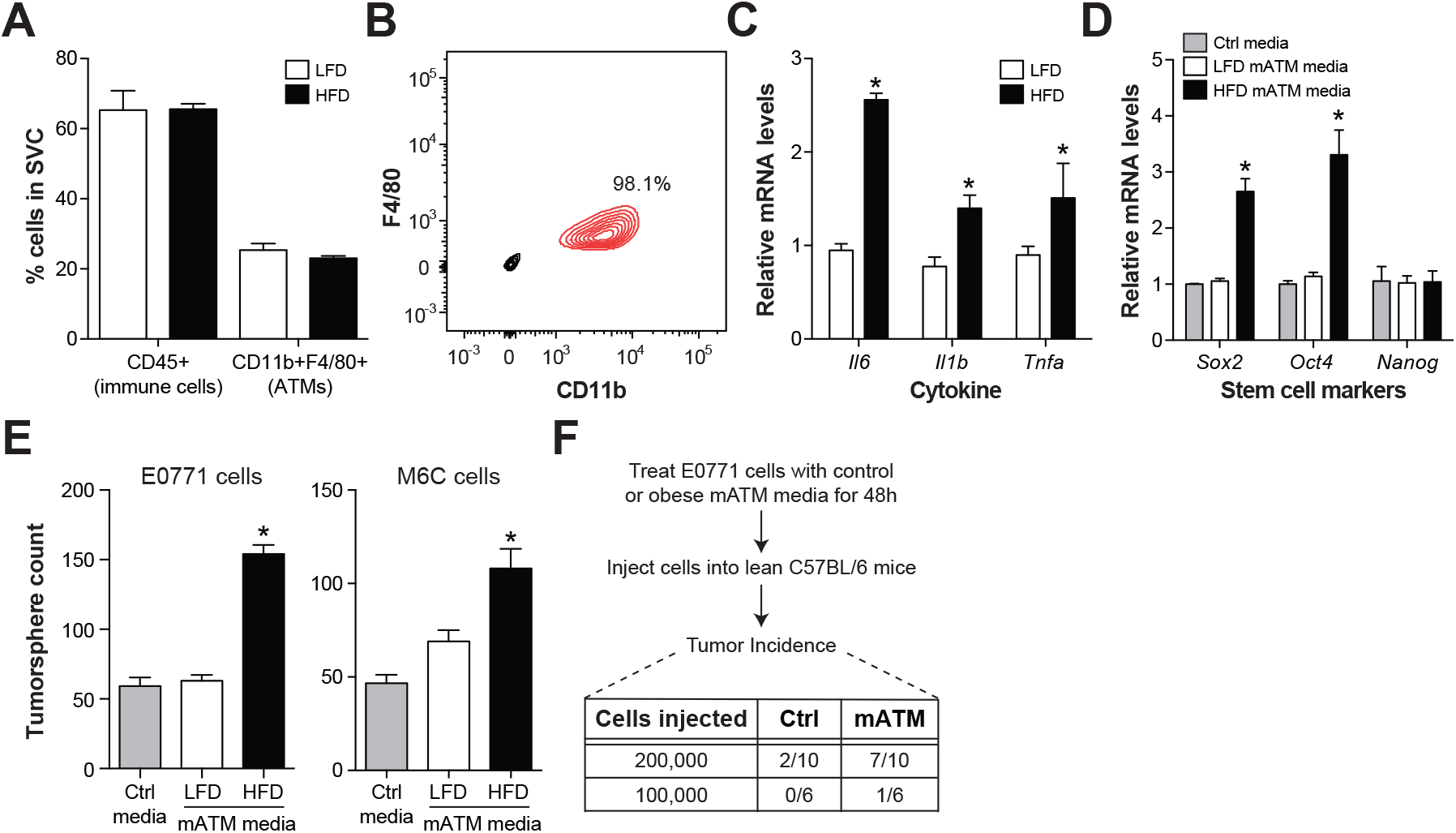
Obese mammary ATMs promote stem-like properties in TNBC cells. Female C57BL/6 mice were fed a LFD or HFD for 12 weeks. *<u>Panel A</u>*: Immune cell composition in the stromal vascular cells (SVC) from mammary adipose tissue. *<u>Panel B</u>*: Purity of mammary ATMs isolated from mice. *<u>Panel C</u>*: Pro-inflammatory cytokine expression in mammary ATMs. *<u>Panels</u> <u>D-F</u>*: TNBC cells were treated with control media (Ctrl) or media conditioned by mammary ATMs from lean (LFD) or obese (HFD) mice. *<u>Panel D</u>*: Stem cell marker expression in E0771 cells. *<u>Panel E</u>*: Tumorsphere formation in E0771 and M6C cells. *<u>Panel F</u>*: Tumor incidence following injection of E0771 cells into mammary fat. Results are mean ± SEM. *, *p*<0.05 Student’s *t*-test. n=3-10/group.

To examine the inflammatory status of mammary ATMs, we purified them from lean and obese mice (**Fig. 2B**) and measured inflammatory cytokine expression. We found that mammary ATMs from obese mice had elevated *Tnfa*, *Il1b*, and *Il6* levels in comparison to those from lean mice (**Fig. 2C**). Thus, although DIO did not induce mammary ATM accumulation in mice, it did increase ATM inflammation in mammary fat; the latter observation is similar to what has been widely reported in visceral fat depots (Xu et al., 2003; Lumeng et al., 2007; Kratz et al., 2014).

Next, we determined whether obese mammary ATMs support stem-like properties in TNBC cells. To explore this, we collected conditioned media from an equal number of mammary ATMs from lean and obese mice and determined whether this media could *i*) induce the expression of the stem cell markers *Sox2*, *Oct4*, and *Nanog* in TNBC cells, and *ii*) promote TNBC cell tumorsphere formation *in vitro* and tumor-initiating potential *in vivo*, functional assays associated with cancer stem cells (Lee et al., 2016).

Whereas treatment with conditioned media from lean mammary ATMs had no effect on stem cell marker expression in E0771 cells, conditioned media from obese mammary ATMs increased the expression of the stem cell markers *Sox2* and *Oct4* (**Fig. 2D**). Obese mammary ATM media also induced tumorsphere formation of E0771 and M6C cells (derived from C3(1)- TAg mice), but lean mammary ATM media could not (**Fig. 2E, S3**). Moreover, pre-treating E0771 cells with obese mammary ATM media increased their tumor-initiating potential *in vivo* (**Fig. 2F**). Together, these findings suggest that obese mammary ATMs are inflamed and promote stem-like properties in TNBC cells, consistent with a pro-tumorigenic phenotype.

### Obese mammary ATMs adopt a metabolically activated (MMe) phenotype

Next, we explored the nature of the pro-inflammatory mammary ATM phenotype in obese mice. Although we previously showed that MMe macrophages are present in visceral and subcutaneous adipose tissue depots in obese humans and mice (Kratz et al., 2014), their presence in mammary adipose tissue has not been investigated. This is important because obesity can elicit substantially diverse effects on different adipose tissue depots (Wang et al., 2013).

We used two approaches to determine whether obesity induced an MMe-like phenotype in mammary ATMs. First, we compared mammary ATMs from lean and obese female C57BL/6 mice for the mRNA expression of markers diagnostic of the M1 (*Cd40*, *Cd38*) and MMe (*Cd36*, *Plin2*) phenotypes (Kratz et al., 2014). We found that obesity induced the expression of MMe markers, but not M1 markers, in mammary ATMs (**Fig. 3A**). These findings suggest that the milder DIO that develops in female mice (Dorfman et al., 2017) is sufficient to support an MMe phenotype in ATMs, and that MMe, rather than M1, is the dominant pro-inflammatory macrophage phenotype in mammary adipose tissue of obese mice.

**Fig. 3.**
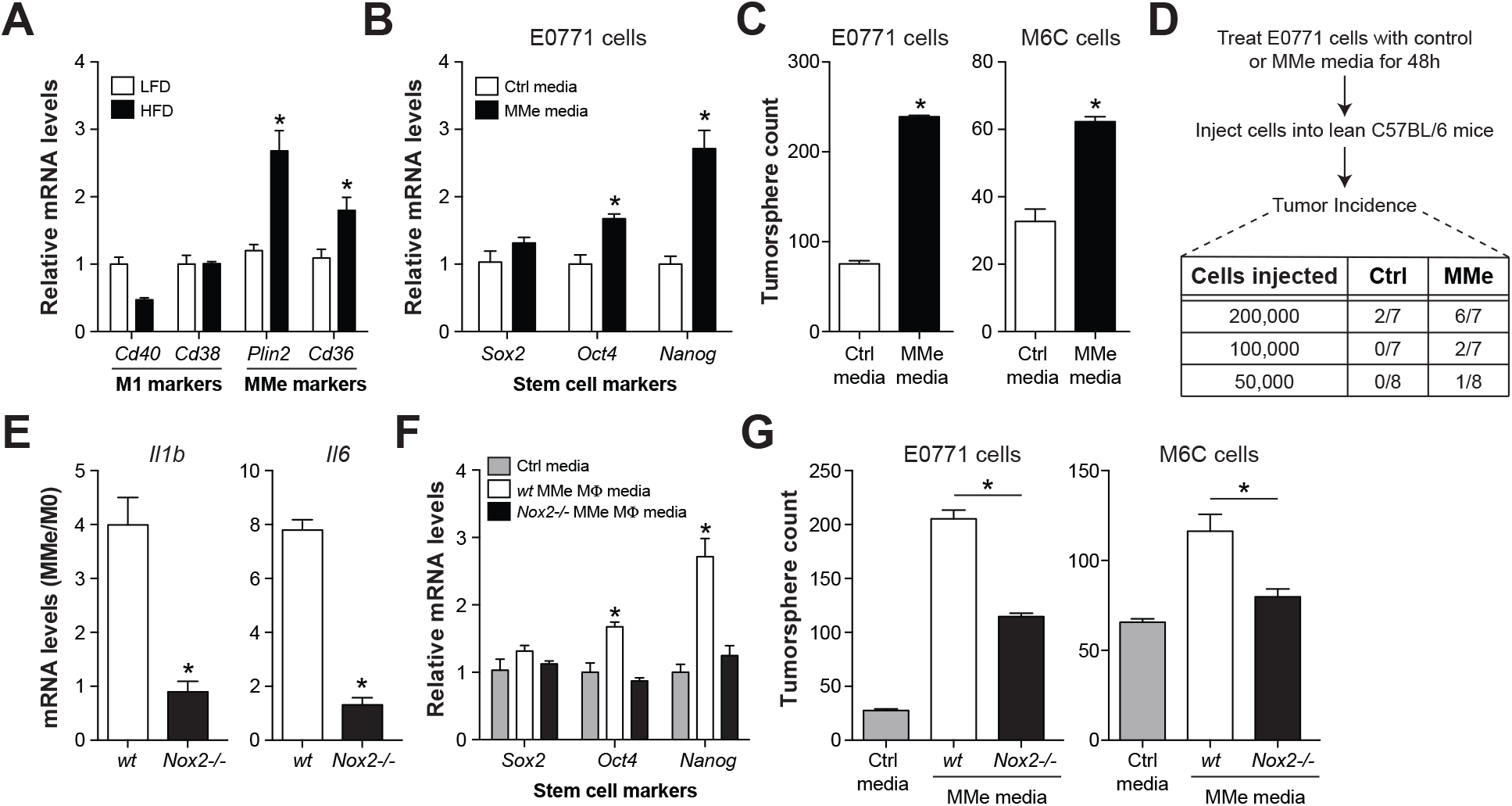
Obese mammary ATMs adopt a metabolically activated (MMe) phenotype. *<u>Panel A</u>*: M1 and MMe marker expression in mammary ATMs from lean and obese C57BL/6 mice. *<u>Panels B-D</u>*: BMDMs were metabolically activated (MMe) *in vitro*. Effect of MMe macrophage media on TNBC stem cell marker expression (*<u>Panel B</u>*), tumorsphere formation (*<u>Panel C</u>*), and tumor incidence following injection of E0771 cells into mammary fat (*<u>Panel D</u>*). *<u>Panels E-F</u>*: *Wt* and *Nox2-/-* BMDMs were metabolically activated. *<u>Panel E</u>*: Cytokine expression in MMe macrophages. *<u>Panels F-G</u>*: Effect of control media and MMe macrophage media on TNBC stem cell marker expression (*<u>Panel F</u>*), and tumorsphere formation (*<u>Panel G</u>*). Results are mean ± SEM. *, *p*<0.05 Student’s *t*-test. n=3-8/group.

Second, we determined whether *in vitro*-derived MMe macrophages could mimic the stem cell promoting properties of obese mammary ATMs. We isolated bone marrow-derived macrophages (M0), metabolically activated them with a mixture of palmitate, insulin, and glucose (MMe), and collected their conditioned media. Conditioned media from MMe macrophages induced the expression of stem-like markers (*Oct4*, *Nanog*) in E0771 cells (**Fig.3B**), tumorsphere formation in E0771 and M6C cells (**Fig. 3C**), and tumor-initiating potential of E0771 cells *in vivo* (**Fig. 3D**).

Together, these findings show that obese mammary ATMs express markers of MMe rather than M1 macrophages, and that *in vitro*-derived MMe macrophages mimic their effects on TNBC stem-like properties.

### Metabolically activated mammary ATMs contribute to the enhanced TNBC tumor formation in obese mice

To determine whether metabolically activated ATMs contribute to TNBC tumorigenesis during obesity, we took advantage of our previous work, which showed that NADPH oxidase 2 (NOX2) is required for the induction of pro-inflammatory cytokine expression in MMe macrophages, but not M1 macrophages (Coats et al., 2017).

We first determined whether deleting NOX2 (*Cybb-/-*) in *in vitro*-derived MMe macrophages could diminish their ability to promote TNBC stem-like properties. As previously described, deleting *Nox2* lowered inflammatory cytokine expression in MMe macrophages (**Fig. 3E**). Moreover, deleting *Nox2* in MMe macrophages attenuated their ability to induce stem-like marker expression and tumorsphere formation in E0771 cells *in vitro* (**Figs. 3F-G, Fig.)**. Thus, the pro-inflammatory and stem cell promoting properties of *in vitro*-derived MMe macrophages are dependent on *Nox2*.

Next, we determined whether NOX2 was required for MMe macrophages to promote tumor formation during obesity. We fed myeloid cell-specific *Nox2*-deficient mice (m*Nox2-/-*) and littermate controls (*wt*) a HFD for 10 weeks. Deleting myeloid cell *Nox2* in female mice did not appreciably impact the metabolic phenotype following 10 weeks of HFD. As previously described in male m*Nox2-/-* mice (Coats et al., 2017), female m*Nox2-/-* mice fed a HFD gained more body weight and adipose tissue mass than *wt* mice, but fasting glucose and insulin levels were unaffected (**Figs. 4A-D**). Mammary and visceral adipose tissue health and liver fat accumulation were also unaffected (**Fig. S4**), which is important because, at least in males, m*Nox2-/-* mice develop late-onset visceral lipoatrophy and hepatosteatosis after 16 weeks of HFD (Coats et al., 2017).

**Fig. 4.**
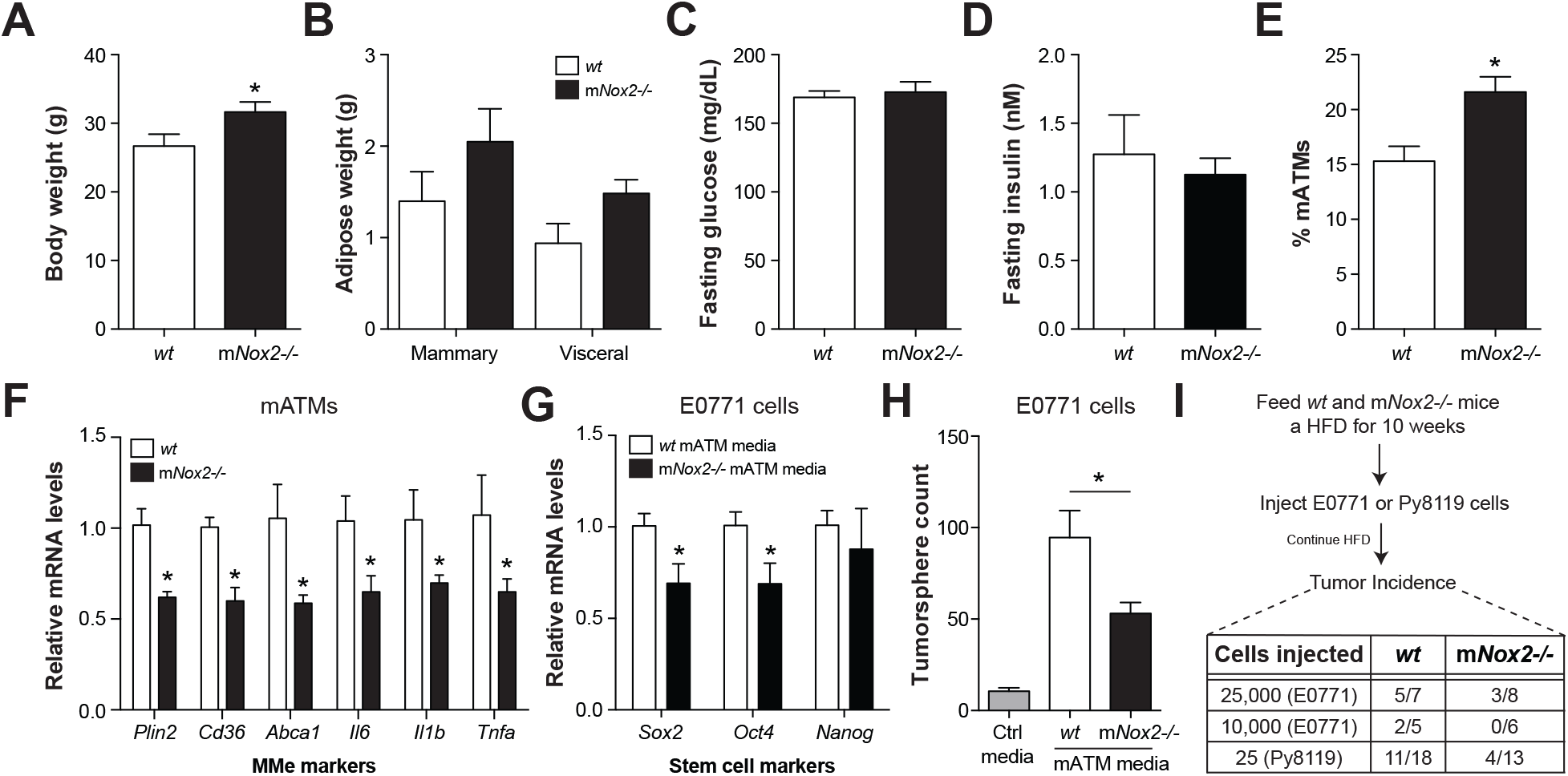
Metabolically activated ATMs contribute to the enhanced TNBC cell tumor formation during obesity. Myeloid cell-specific LysM-cre^+/-^ *Nox2^fl/fl^* mice (m*Nox2-/-*) and littermate control *Nox2^fl/fl^* mice (*wt*) were fed a HFD for 10 weeks. *<u>Panel A</u>*: Body weight. *<u>Panel</u> <u>B</u>*: Mammary and visceral adipose tissue weight. *<u>Panels C-D</u>*: Fasting glucose and insulin levels. *<u>Panel E</u>*: Percent ATMs in mammary adipose tissue. *<u>Panel F</u>*: MMe marker expression in mammary ATMs. *<u>Panels G-H</u>*: Effects of mammary ATM media on stem cell markers (*<u>Panel G</u>*), and tumorsphere formation (*<u>Panel H</u>*) in E0771 cells. *<u>Panel I</u>*: Tumor incidence following injection of E0771 or Py8119 cells into mammary fat. Results are mean ± SEM. *, *p*<0.05 Student’s *t*-test. n=3-15/group.

Although the percentage of mammary ATMs was elevated in m*Nox2-/-* mice (**Fig. 4E**), analysis of these ATMs revealed attenuated expression of all MMe markers tested, including *Tnfa*, *Il6*, *Il1b*, *Abca1*, *Cd36*, and *Plin2* (**Fig. 4F**). Thus, these female m*Nox2-/-* mice afford an opportunity to study the role of MMe macrophages in TNBC tumor formation during obesity in the absence of substantive changes to the overall metabolic phenotype.

We used two approaches to determine the impact of deleting *Nox2* in myeloid cells on TNBC tumor formation during obesity. First, we isolated mammary ATMs from obese *wt* or m*Nox2*-/- mice and measured their capability to promote TNBC stem-like properties. Mammary ATMs from obese m*Nox2-/-* mice had decreased ability to promote stem cell marker expression (*Sox2*, *Oct4*) and tumorsphere formation of E0771 cells (**Figs. 4G-H, S3**).

Second, we used a limiting dilution assay to investigate whether deleting *Nox2* from myeloid cells could attenuate the ability of DIO to promote the tumor-initiating potential of TNBC cells *in vivo*. Deleting *Nox2* from myeloid cells decreased tumor incidence in obese mice injected with E0771 or Py8119 cells (**Fig. 4I**). This decreased tumor incidence was not absolute, which may be explained by *i*) the slight increases in body weight, mammary fat mass, and mammary ATM number in m*Nox2-/-* mice (see Figs. 4A, 4E), and *ii*) the inability of *Nox2* deletion to completely block the induction of stem-like properties in TNBC cells (see Figs. 3G, 4H).

Collectively, these data suggest that *Nox2* is required for the metabolic activation of mammary ATMs *in vivo*, and highlight the contribution of MMe macrophages in mammary fat to the increased TNBC tumor formation during obesity.

### Metabolically activated mammary ATMs release cytokines that signal through GP130 on TNBC cells to promote their stem-like properties

Previous studies showed that pro-inflammatory cytokines bind glycoprotein 130 (GP130) to induce STAT3 phosphorylation, which in turn, promotes the stem-like properties of many types of cancer cells (Korkaya et al., 2011). One well-studied GP130 ligand is IL6, whose expression is elevated in metabolically activated mammary ATMs from obese mice, and attenuated by deleting *Nox2* (see Figs. 2C, 4F). Because IL11, OSM, LIF, CNTF, and CTF1 can also signal through GP130 (Silver and Hunter, 2010), we examined their regulation in mammary ATMs, and found that obesity increased their transcript expression (**Fig. 5A**). Moreover, treating E0771 or M6C cells with mammary ATM conditioned media from obese relative to lean mice induced STAT3 phosphorylation (**Fig. 5B, 5D**), a key effector of GP130 signaling. *In vitro*-derived MMe macrophages similarly up-regulated the expression of all GP130 ligands tested and their media also induced STAT3 phosphorylation in E0771, M6C, and Py8119 cells (**Fig. S5**).

**Fig. 5.**
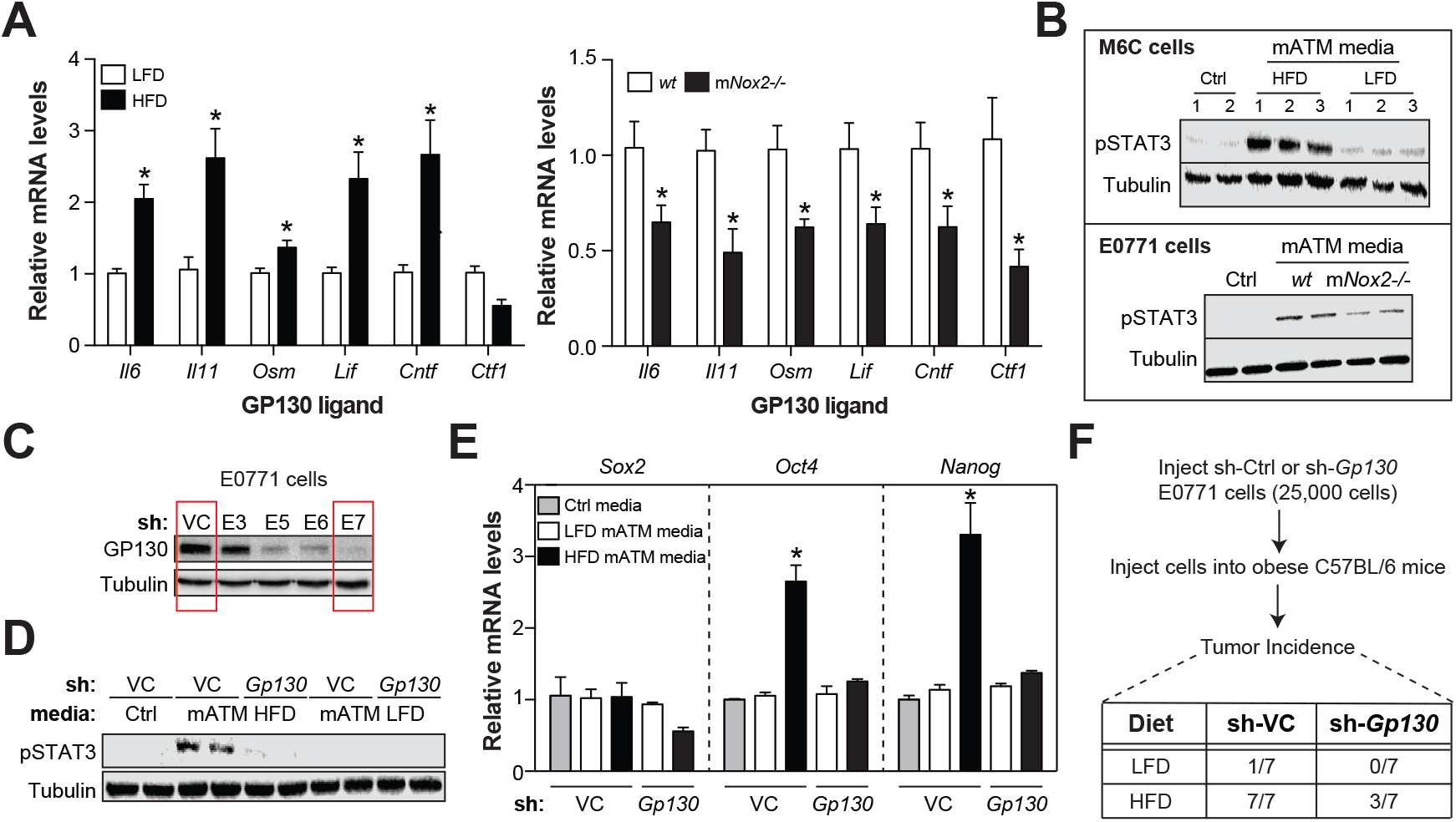
MMe macrophages release cytokines that signal through GP130 to drive TNBC cell stemness. *<u>Panel</u>* A: GP130 ligand expression in mammary ATMs isolated from lean and obese C57BL/6 mice, or obese *mNox2-/-* mice and *Nox2^fl/fl^* mice (*wt*) littermate controls. *<u>Panel B</u>*: Effects of mammary ATM media on STAT3 phosphorylation in TNBC cells. *<u>Panel C</u>*: E0771 cells were treated with vector control shRNA (VC) or *Gp130* shRNA (E3, E5, E6, E7) and GP130 knockdown was confirmed by western blotting. *<u>Panels D-E</u>*: Effect of GP130 knockdown (E7) on the ability of mammary ATM media to induce STAT3 phosphorylation (*<u>Panel D</u>*) and stem cell marker expression (*<u>Panel E</u>*) in E0771 cells. *<u>Panel F</u>*: Tumor incidence following injection of sh-VC or sh-Gp130 (E7) E0771 cells into mammary fat of lean or obese C57BL/6 mice. Results are mean ± SEM. *, *p*<0.05 Student’s *t*-test. n=3-7/group.

We further determined whether GP130 ligand expression was lowered in mammary ATMs from obese m*Nox2-/-* mice, a perturbation that attenuated both the MMe phenotype of mammary ATMs and the ability of obesity to promote tumor formation (see Fig. 4). We found that mammary ATMs from obese *mNox2-/-* mice had decreased GP130 ligand expression (**Fig. 5A**) and diminished capability to induce STAT3 phosphorylation in E0771 cells (**Fig. 5B**), relative to mammary ATMs from obese *wt* mice.

Based on these findings, we reasoned that GP130 may be required for obese mammary ATMs to promote stem-like properties in TNBC cells. Because each of the GP130 ligands elevated in obese mammary ATMs bind distinct receptors that heterodimerize with GP130 (Silver and Hunter, 2010), we depleted GP130 to broadly attenuate signaling from multiple ligands.

We used shRNA to knockdown GP130 levels in E0771 cells, confirmed knockdown using several clones (**Fig. 5C**), and studied effects of obese mammary ATMs on TNBC cells. We found that knocking down GP130 attenuated the ability of obese mammary ATM media to induce STAT3 phosphorylation and to increase stem cell marker expression (*Oct4*, *Nanog*) in E0771 cells (**Figs. 5D-E**). Knocking down GP130 also lowered the ability of *in vitro*-derived MMe macrophages to induce STAT3 phosphorylation and tumorsphere growth in E0711 cells (**Fig. S5**), suggesting that this pathway may be utilized by MMe macrophage to promote stem-like properties in TNBC cells.

We further explored whether GP130 signaling was required for DIO to promote the tumor-initiation potential of TNBC cells *in vivo* by injecting sh-control or sh-*Gp130* E0771 cells into leand and obese female mice and monitoring tumor incidence. *Gp130* knockdown in E0771 cells attenuated the ability of obesity to promote tumor formation *in vivo* (**Fig. 5F**), and this decrease was comparable to the effect observed in obese m*Nox2-/-* mice (see Fig. 4I).

### Human MMe macrophages are present in mammary adipose tissue of obese women

Our findings suggest that DIO promotes an MMe phenotype in mammary ATMs, which overexpress cytokines that signal through GP130 to promote the stem-like properties of TNBC cells in mice. We investigated whether this mechanism may also be operative in humans.

We determined whether MMe macrophages were increased in mammary adipose tissue of obese women (BMI > 30kg/m^2^) in comparison to non-obese women (BMI < 30kg/m^2^) undergoing mammary reductions. To this end, we isolated the SVC from mammary adipose tissue and studied ATMs (defined as CD45^+^CD11b^+^CD14^+^) by flow cytometry.

The number of ATMs (defined as CD45^+^CD11b^+^CD14^+^) in the SVC from mammary fat was significantly elevated in obese women relative to non-obese women (**Fig. 6A-B, Fig. S6**). To explore the activation status of mammary ATMs, we stained them with antibodies against M1 markers (CD319 and CD38) and MMe markers (ABCA1 and CD36). These analyses showed that MMe ATMs, and not M1 ATMs, accumulate in mammary adipose tissue during obesity (**Fig. 6A-B**). Indeed, the percent of mammary ATMs expressing MMe markers was significantly and positively correlated with BMI (*p*<0.001, R^2^ = 0.67), while the percent M1 ATMs was not correlated (*p*=0.14, R^2^ = 0.14) (**Fig. 6C**).

**Fig. 6.**
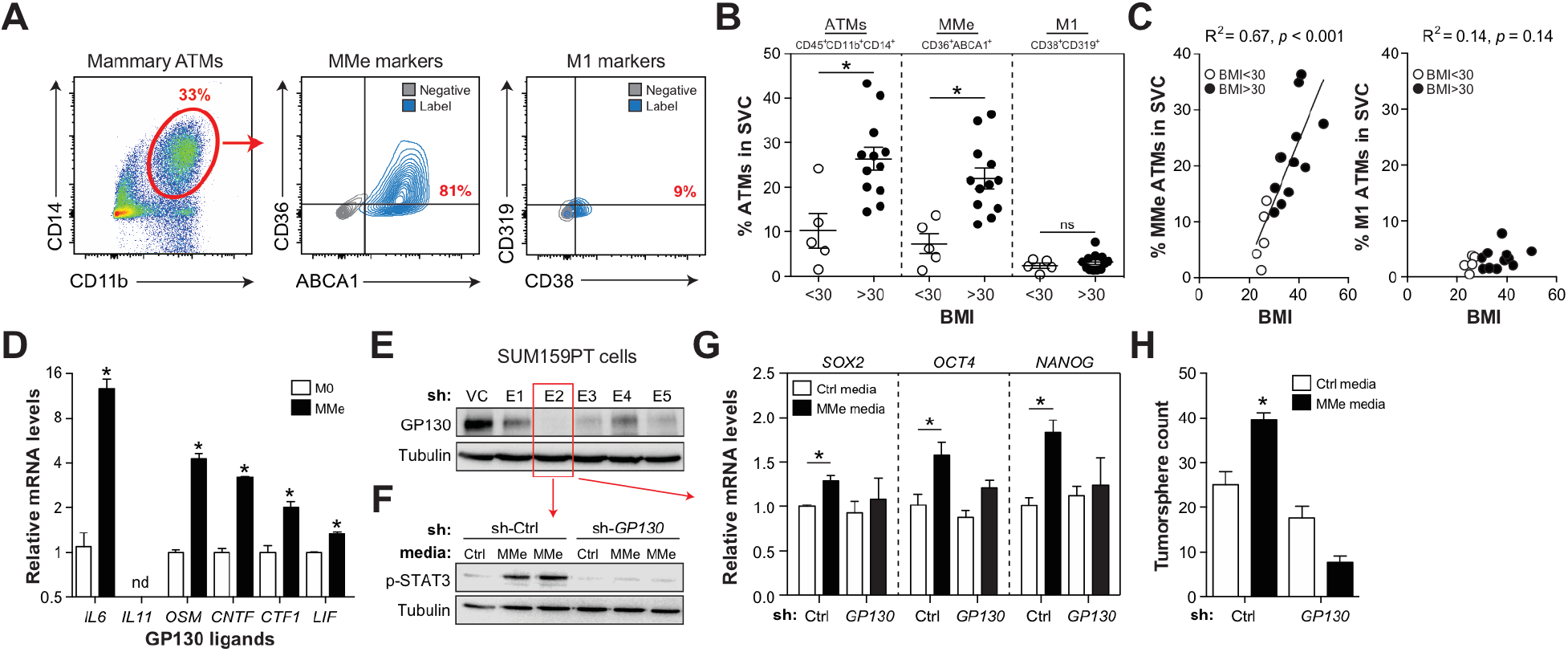
MMe macrophages are present in mammary fat of obese women, and promote stem-like properties in TNBC cells. *<u>Panels A-C</u>*: Mammary adipose tissue was obtained from non-obese (BMI<30, n=5) and obese (BMI>30, n=12) women. *<u>Panel A</u>*: Flow cytometric analysis of mammary ATMs. One subject is shown as an example. *<u>Panel B</u>*: Percent ATMs, MMe ATMs, and M1 ATMs in the stromal vascular cells (SVC) obtained from mammary fat. *<u>Panel C</u>*: Linear regression analysis of percent MMe and M1 ATMs in mammary fat with BMI. *<u>Panel D</u>*: GP130 ligand expression in unstimulated (M0) and metabolically activated (MMe) HMDMs; nd = not detected. *<u>Panel E</u>*: Human SUM159PT cells were treated with control shRNA (VC) or *GP130* shRNA (E1-E5) and GP130 knockdown was confirmed by western blotting. *<u>Panels F-H</u>*: Effect of *GP130* knockdown (E2) on the ability of human MMe media to induce STAT3 phosphorylation (*<u>Panel F</u>*), stem cell marker expression (*<u>Panel G</u>*), and tumorsphere formation (*<u>Panel H</u>*) in SUM159PT cells. Results are mean ± SEM. *, *p*<0.05 Student’s *t*-test. n=3- 12/group.

Interestingly, the majority of mammary ATMs from non-obese and obese subjects stained positive for MMe markers (**Fig. S6**), suggesting that obesity did not induce a phenotypic switch in ATMs towards the MMe phenotype. Instead, we observed an increase in the number of MMe macrophages in the SVC of obese mammary fat (see Figs. 6B-C). These data are in contrast to what we observed in mammary adipose tissue in mice, where obesity induced MMe marker expression in ATMs without altering their number (see Fig. 2). Thus, although the mechanisms are different between mice and humans, the common endpoint is an increased presence of MMe macrophages in mammary adipose tissue during obesity.

Next, we determined whether human MMe macrophages could signal through GP130 to promote human TNBC cell stem-like properties. Because we were unable to isolate enough ATMs from human mammary adipose tissue, we tested this hypothesis using human monocyte-derived macrophages (HMDMs) exposed to MMe polarizing stimuli *in vitro*.

*In vitro*-derived human MMe macrophages over-expressed all GP130 ligands tested, including *IL6*, *OSM*, *CNTF*, *CTF1*, and *LIF* (**Fig. 6D**). Moreover, human MMe macrophage media induced STAT3 phosphorylation, stem cell marker expression (*SOX2*, *OCT4*, *NANOG*), and tumorsphere growth of SUM159PT cells (**Figs. 6E-H**), a human TNBC cell line (Neve et al., 2006). Importantly, all of these effects were diminished when GP130 was knocked down in SUM159 cells by shRNA (**Figs. 6E-H**).

Collectively, these findings suggest that during obesity, MMe macrophages accumulate in mammary adipose tissue, and that human MMe macrophages, like their mouse counterparts, can promote stem-like properties in TNBC cells through GP130 signaling.

## DISCUSSION

Our findings support a model wherein DIO induces an MMe phenotype in mammary ATMs, which overexpress cytokines that signal through GP130 to induce stem-like properties in TNBC cells. Three lines of evidence support the idea that metabolically activated mammary ATMs promote TNBC tumor formation during obesity. First, we show that obesity induces human and murine mammary ATMs to express markers of the MMe phenotype. Second, we demonstrate that *in vitro*-derived murine and human MMe macrophages mimic the effects of obese mammary ATMs on stem-like properties of TNBC cells, both in terms of mechanism and function. Third, we show that ablating *Nox2*, a gene required for MMe macrophage polarization, attenuated the ability of obese mammary ATMs to promote TNBC stemness *in vitro*, and the ability of obesity to promote TNBC tumor formation *in vivo*. The importance of this mechanism is underscored by our demonstration that it is utilized by *in vitro* and *in vivo*-derived MMe macrophages to induce stem-like properties in multiple TNBC cell lines of murine and human origin.

These findings, together with our previous work (Kratz et al., 2014), suggest that the MMe phenotype may be broadly applicable to ATMs in multiple adipose tissue depots during obesity in humans and mice, and that these cells perform a wide array of functions that impact the metabolic phenotype and its associated co-morbidities. We previously showed that MMe-like ATMs in visceral fat over-express inflammatory cytokines to exacerbate insulin resistance during early obesity, but also protect against insulin resistance in prolonged obesity by clearing dead adipocytes through a lysosomal exocytosis pathway (Coats et al., 2017). Our new findings suggest that MMe-like ATMs in mammary fat overexpress GP130 ligands that promote TNBC stem-like properties, and it is possible that a similar MMe macrophage mechanism may promote ER+ breast cancer and other obesity-associated cancers arising in adipose tissue rich environments (eg. ovarian, colon, brain). However, deciphering the contribution of MMe macrophages in ER+ breast cancer may be challenging because obesity is also known to promote ER+ breast cancer by increasing local production of estrogen in adipose tissue (Subbaramaiah et al., 2011).

Cancer cells with stem-like properties are known to promote tumor initiation and metastasis (Grivennikov et al., 2010). These stem-like properties may be activated by mutation of key genes or epigenetic regulators in cancer cells, or by signals from the tumor microenvironment (Karnoub et al., 2007; Liu et al., 2011). Indeed, inflammatory cytokines (such as IL6) arising from tumor cells or an altered tumor microenvironment can signal through GP130 to induce JAK/STAT signaling and drive cancer cell stemness (Korkaya et al., 2011; Lu et al., 2014). Constitutively activated STAT3 has been implicated in the initiation of many types of cancer, including breast cancer (Ling and Arlinghaus, 2005), as well as the promotion of invasion, migration, epithelial-to-mesenchymal transition, and cancer stem cell self-renewal and differentiation (Korkaya et al., 2011; Yuan et al., 2015). Our findings suggest that the chronic metabolic activation of ATMs in mammary fat promotes TNBC tumor formation through this mechanism and could contribute to metastatic progression as well.

Importantly, disabling this MMe-GP130-stemness pathway (by deleting *Nox2* in macrophages or attenuating *Gp130* in TNBC cells) did not completely block increased tumor formation during obesity. It is therefore likely that additional DIO-induced factors can contribute to this phenotype. For example, leptin, an adipocyte-derived hormone that is increased during obesity, has been shown to promote cancer cell stemness in breast cancer (Chang et al., 2015). Free fatty acids (FFAs) can also promote stem-like properties in cancer cells (Ye et al., 2016), and the increased adipose tissue mass and adipocyte insulin resistance would increase FFA levels in mammary adipose tissue during obesity. These increased FFA levels would also stimulate MMe polarization in mammary ATMs, which our findings suggest substantially contributes to TNBC cell stemness during obesity.

More generally, our findings reinforce the idea that tumorigenesis is regulated both by intrinsic properties of cancer cells (ie. genetic alterations) as well as the tissue-specific niche in which the tumor develops (Polyak et al., 2009). From this perspective, obesity may be conceptualized as a pathological state that facilitates tumorigenesis by creating tumor permissive conditions in multiple tissues. For example, our studies demonstrated that obesity-induced ATM inflammation in mammary fat promotes TNBC tumor formation, while previous studies showed that obesity-induced neutropenia in the airway facilitates TNBC tumor metastasis (Quail et al., 2017). Although the specific mechanisms underlying these observations are distinct, they can be integrated through obesity’s ability to induce chronic inflammation. Indeed, chronic inflammation, from a multitude of underlying sources (eg. ulcerative colitis, Hepatitis C, pancreatitis), is known to promote cancer cell stemness and increase the risk of many types of cancers (Multhoff et al., 2011).

The mechanism by which MMe macrophages induce stem-like phenotypes in tumor cells reveals a number of potential therapeutic approaches to counteract the effects of obesity. NOX2 inhibitors such as gp91sd-tat (Smith et al., 2015) may be an attractive approach for attenuating MMe macrophage inflammation in obesity-driven TNBC. GP130 inhibitors such as Bazedoxifene (Xu and Neamati, 2013), may also be useful for blocking the ability of MMe macrophage-derived cytokines to induce TNBC cell stemness. However, this approach may require the development of new GP130 inhibitors because MMe macrophages overexpress many GP130 ligands and Bazedoxifene’s ability to inhibit the GP130/STAT3 pathway is limited to IL6 and IL11 (Wu et al., 2016). Finally, JAK1/2 inhibitors may have therapeutic value in blocking signaling downstream of MMe-derived cytokine interactions with GP130, and the JAK1/2 inhibitor Ruxolitinib is currently in clinical trials for treatment of TNBC (Harrison et al., 2012). Future studies will be needed to test the efficacy of these therapeutics in treating obesity-driven TNBC.

## EXPERIMENTAL PROCEDURES

### Regulatory

Animal studies were approved by the University of Chicago IACUC (ACUP #72209 and ACUP #72228). Human studies were approved by the Institutional Review Boards at the University of Chicago (IRB16-0321) and Northwestern University (NU 11B04).

### Subject recruitment

Human breast adipose tissue was obtained from women undergoing breast reduction surgery surgeries at Northwestern Memorial Hospital. Exclusion criteria included cancer or any other breast related diseases. Human breast adipose tissue was collected by surgeons and tissue was processed immediately to obtain the stromal vascular cells (SVC) for flow cytometric analyses (see below).

### Mice

Wild-type and *Nox2-/-* (*Cybb-/-*) female mice on the C57BL/6 background, and C3(1)- TAg mice are from Jackson Labs. Myeloid cell specific *Nox2-/-* mice (m*Nox2-/-*) were generated by crossing *Cybb^fl/fl^* mice (Sag et al., 2017) with *LysM-cre* knock in mice (Jackson Labs, 004781) to generate LysM-cre^+/-^ *Cybb^fl/fl^* and litter mate control *Cybb^fl/fl^* mice as previously described (Coats et al., 2017). Mouse genotype was confirmed by PCR (see **Table S1** for primers).

### DIO studies

Female C57BL/6 or C3(1)-TAg female mice were placed on a low-fat (Harlan) or 60% high-fat diet (D12451, Research Diets Inc.) at 6 weeks of age for up to 12 weeks. Body weight was monitored every week.

### Plasma metabolic measurements

Mice were fasted for 3h and blood glucose levels were measured with a One Touch Ultra 2 glucometer (Lifescan) and serum insulin levels were measured by ELISA (Millipore).

### TNBC cell lines

The TNBC cells lines used in this study include: E0771 cells, originally isolated from a spontaneous breast adenocarcinoma in C57BL/6 mice (Casey et al., 1951); M6C cells, originally isolated from the C3(1)/SV40 Large T-antigen transgenic mouse model (Holzer et al., 2003); Py8119 cells, derived from a mammary adenocarcinoma that spontaneously arose in a MMTV-PyMT transgenic C57BL/6 female mouse (Gibby et al., 2012); and SUM159PT cells, derived from a primary TNBC tumor from a patient with invasive ductal carcinoma (Neve et al., 2006). Py8119 and SUM159PT cells were obtained from ATTC and cultured according to their guidelines. All other cells were maintained in DMEM supplemented with 10% FCS.

### Cancer-cell isolation from tumor

Tumors were digested with collagenase and hyaluronidase (1:3) and 0.1 mg/mL DNase I (Worthington, Lakewood, NJ), filtered through 70-μm mesh, incubated with RBC lysis buffer, filtered through 40-μm mesh, and resuspended in PBS with 1% BSA. To isolate cancer cells for qRT-PCR analysis, mononuclear cells were depleted by centrifuging using Ficoll-Paque PREMIUM, with a density of 1.077± 0.001 g/mL for 40 minutes at 400 g after RBC lysis. The pellet at the bottom was re-suspended in RLT buffer for RNA isolation.

### ATM isolation and analysis

SVC was obtained by digesting human or mouse mammary adipose tissue with collagenase type 2 for 60 minutes and filtering the supernatant with 40 μm filter after red blood cell lysis. Mammary ATMs in the SVC obtained from murine and human mammary fat were interrogated by flow cytometry (see **Figs. S2,S6** for the gating strategy). Murine mammary ATMs were isolated using anti-CD11b antibody coupled to magnetic beads as previously described (Kratz et al., 2014) and purity was assessed by flow cytometry. ATMs were interrogated by qRT-PCR and media was collected for functional assays.

### Differentiation and metabolic activation of murine bone marrow-derived macrophages and human monocyte-derived macrophages

Murine bone marrow-derived macrophages (BMDMs) were generated and metabolically activated as previously described (Kratz et al., 2014). Briefly, bone marrow was flushed from the bones of the hind legs and differentiated to macrophages by culturing for 6 days in six-well plate in DMEM with 10% FCS plus conditioned media from L929 cells at a 1:3 ratio. Human peripheral blood monocytes were isolated from healthy donors and differentiated to macrophages in the presence of M-CSF (125 ng/mL). For MMe activation, macrophages were treated with glucose (30mM), insulin (10nM), and palmitate (0.3mM) for 24h. Conditioned media (CM) was generated by culturing primary immune cells in regular DMEM serum free media for 24hrs, using .6 mL of media for every 1 million macrophages seeded.

### Limiting dilution assays

E0771 or PY8119 cells were transplanted into the #4 mammary fat pad. Tumors were considered established when they became palpable for 2 consecutive weeks. Analyses of tumor incidence were confirmed by two independent investigators. To test the effects of ATMs or MMe macrophages on E0771 tumor-initiating potential, we serum starved the cells overnight, treated them with macrophage conditioned media for 24h in DMEM supplemented with 2.5% FBS, counted the cells, and injected them into the mammary fat pad.

### Tumorsphere assays

E0771, M6C, or SUM159PT cells were plated at 500 cells/well of a ultra-low attachment 12-well plate (Corning) in standard mammosphere media comprising DMEM-F12 supplemented with FGF (20ng/mL, Gold Bio.), EGF (20ng/mL, Gold Bio.), Heparin (4μg/mL, Sigma) and B27 supplement (Life Technologies). Tumorspheres were counted following 5-8 days of growth. To study the effects of macrophage media on tumorsphere formation, tumor cells were pre-treated with macrophage conditioned media for 48h in DMEM supplemented with 2.5% serum. At this time, cells were collected, counted, and plated for tumorsphere assays. To study the effects of macrophage media on tumorsphere formation, tumor cells were serum-starved overnight and pre-treated with macrophage conditioned media for 24h in DMEM supplemented with 2.5% serum.

### GP130 knockdown

Short hairpin RNA (shRNA) specific for murine or human *Gp130* ligated into the lentiviral vector pLKO-1 were purchased from Dhramacon. Virus particles were packaged, E0771 and SUM159PT cells were infected, and infected cells were selected for by treatment with puromycin. Lentiviral pLKO-1 vector without shRNA was used as a control. Knockdown of GP130 was verified by immunoblotting.

### Antibodies

Antibodies for flow cytometric measurements of CD45, CD11b, CD206, CD14, ABCA1, CD36, CD38, CD45, and CD206, CD319, F480 were purchased from BD Biosciences (San Jose), Beckman Coulter (Danvers), Novus Biologicals (Littleton) BioLegend (San Diego), eBioscience (San Diego), or Miltyeni (Auburn).

### Statistics

Statistical significance was assessed using an unpaired, two-tailed, Student’s *t*-test. Replicate numbers for each experiment are indicated in the figure legends.

### qRT-PCR

RNA was isolated using Qiagen Midi-prep kits, reverse transcribed with Quantiscript (Qiagen) using random hexamers (Invitrogen), and mRNA levels were measured with specific primers (**Table S1**) using SYBR green on a One Step Plus system (Applied Biosystems). Relative levels of each target gene were calculated using the ∆∆Ct formula, using 18S RNA as a control.

## ACKNOWLEDGEMENTS

This research was supported by grants from the National Institutes of Health (R01DK102960 (L.B.), R01CA184494 (M.R.R.), F99CA212477 (support for P.T.), the Cancer Research Foundation (L.B.), The Avon Foundation (S.A.K.), the Bernice Goldblatt Endowment Fellowship, University of Chicago (support for C.C.), and the Department of Health via a National Institute for Health Research Biomedical Research Centre award to Guy’s & St Thomas’ NHS Foundation Trust and King’s College London (support for A.M.K.). We would like to thank Drs. Ramesh Ganju (Ohio State University) and Jeffrey Green (National Cancer Institute) for providing the E0771 and M6C cell lines respectively.

## AUTHOR CONTRIBUTIONS

Conceptualization: L.B., M.R.R, P.T. Investigation: P.T., A.B., C.C., G.Z., K.Q.S., Y.X. Writing – Original Draft: L.B., M.R.R., P.T. Writing – Reviewing & Editing: All authors. Supervision: L.B., M.R.R., S.A.K. Funding Acquisition: L.B., M.R.R., S.A.K.

